# Disentangling sex-ratio meiotic drive in *Drosophila* a century after its original discovery

**DOI:** 10.1101/2025.05.14.653954

**Authors:** Takehiro K. Katoh, Yinjia Chen, Naohito Takatori, Masafumi Nozawa

## Abstract

Sex ratio is distorted when selfish elements linked to the sex chromosome cause meiotic drive. Since the first report on the drive in *Drosophila obscura* in 1928, this phenomenon has been reported in diverse taxa. However, the molecular and evolutionary mechanisms of the drive have remained largely elusive, even in *D. obscura*. We therefore reinvestigated this original example by comparing *D. obscura* strains with and without distortion (SR and ST strains, respectively). Crossing experiments indicated the presence of a driver on the X chromosome and predicted the existence of a suppressor. Using comparative genomics, we identified an X-linked gene, *Gcna*, as a candidate driver; the region encoding the SprT-like domain of this gene was massively duplicated only in the SR genome. We further identified a candidate suppressor, *miR-9c*, that interacted with *Gcna* in the SR strain but not with that in the ST strain. *Gcna* with amplified SprT-like domains likely targets transient TOP2-DPC on the Y chromosome and may disrupt or kill sperm harboring the Y chromosome; this was supported by the small number of sperm in the SR strain. Our findings demonstrate an evolutionary arms race between driver and suppressor genes in a longstanding example of sex-ratio meiotic drive.

## Introduction

Chromosomes generally follow balanced Mendelian segregation (i.e., the transmission rate for each of the homologous chromosomes into gametes is ∼50%). However, the Mendelian law of segregation can be violated and one of the most prominent causes of this violation is known as meiotic drive, in which so-called selfish elements elevate their own transmission rate into gametes during meiosis. If such a selfish element is located on a sex chromosome, it distorts not only Mendelian segregation but also the sex ratio. Nearly 100 years ago, Gershenson^1^ first documented this phenomenon in *Drosophila obscura* collected near Moscow, Russia. Some *D. obscura* strains exhibited a female-biased sex ratio, whereas others did not. This sex-ratio distortion was suggested to be caused by a genetic element, or driver, that is located on the X chromosome and kills the Y-chromosome-bearing sperm. Since Gershenson’s report, sex-ratio meiotic drive has been described across diverse multicellular organisms^2,3^.

According to Fisher’s theory of the sex ratio^4^, however, a skewed sex ratio is not evolutionarily stable and potentially leads to population extinction^5^. Consequently, different genetic elements that counteract the effects of drivers, referred to as suppressors, often emerge, with the potential for an arms race between drivers and suppressors^5^. If this process continues in one population but not in the other, the two populations may be reproductively isolated^6^. Therefore, sex-ratio meiotic drive can impact not only sex-chromosome evolution^7^ but also speciation^8^. Moreover, this biological phenomenon has been utilized in applied science; for example, artificial meiotic drive systems have been used for the control of pests, such as mosquitos^9^. Despite extensive research, the drivers that cause sex-ratio distortion and their mode of action have rarely been clarified at the molecular level. In Diptera, for instance, *Dox* (*Distorter on X*) in *D. simulans*, with unknown function, is the only gene that has been identified as a driver of sex-ratio distortion^10^.

In this study, we re-discovered a female-biased sex ratio in a laboratory strain of *D. obscura* (SR strain, hereafter), different from those described by Gershenson^1^ almost a century ago. In addition, another strain with a balanced sex ratio (ST strain, hereafter) was identified in our laboratory. We used traditional genetic experiments and several next-generation sequencing technologies to identify candidate driver and suppressor genes in *D. obscura.* We discuss how sex-ratio distortion and recovery to a balanced sex ratio has occurred at the molecular level under an evolutionary arms race. Our study represents significant progress toward dissecting the original case of sex-ratio distortion after almost 100 years.

## Results

### Sex-ratio distortion was caused by X-linked driver(s) and autosomal or Y-linked suppressor(s) present in the SR strain

We first examined the possibility of endosymbiont-induced sex-ratio distortion. If the observed distortion was caused by endosymbionts, such as *Wolbachia*, a balanced sex ratio would be recovered by antibiotic treatment. However, the SR strain treated with tetracycline for two generations still exhibited a skewed sex ratio and the proportion of males did not differ significantly between the treatment and control conditions (*p* = 0.1767 by Fisher’s exact test, Supplementary Fig. S1). Therefore, the distortion in the SR strain is unlikely to have been caused by cytoplasmic bacterial agents.

To examine how the driver is inherited, we next conducted crossing experiments using the SR and ST strains (Fig. 1a). The crosses using the SR males exhibited a small proportion of male progeny, irrespective of the female strain (crosses B and D in Fig. 1a, b). By contrast, the F_1_ generation derived from ST males (crosses A and C) showed a more or less balanced sex ratio, regardless of the female strain. These results indicated that the driver in the SR strain is located on the X chromosome and drive likely occurs during male meiosis.

**Fig. 1.**
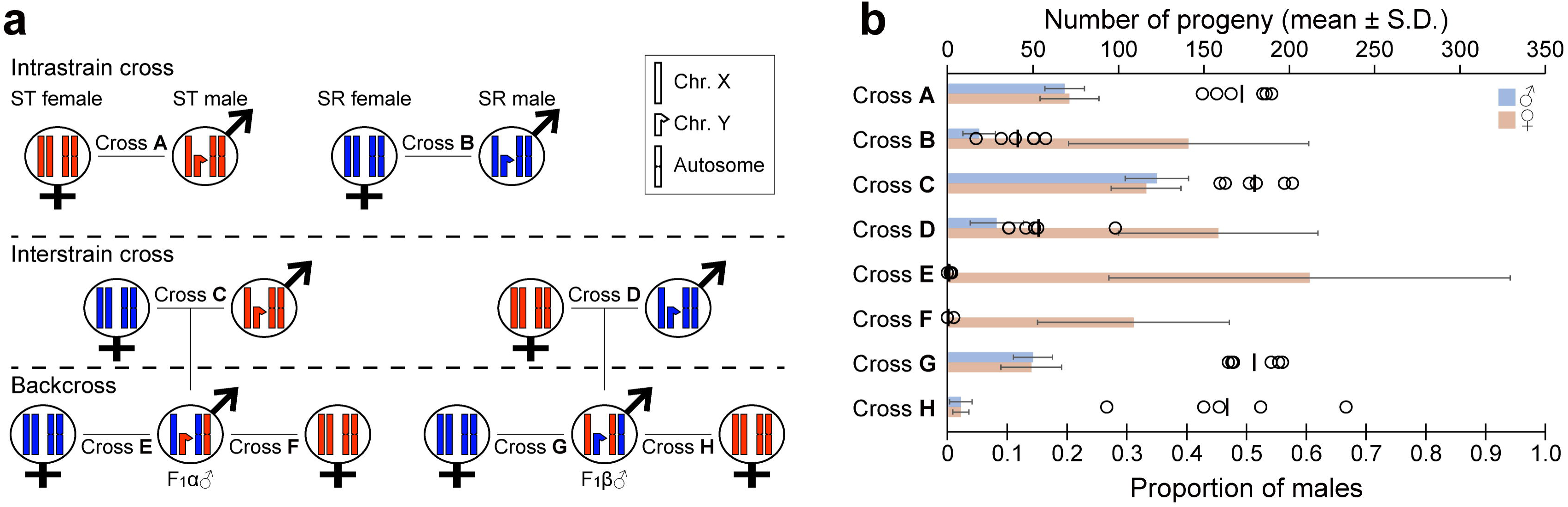
Overview of cross experiments using the SR and ST strains. **a,** Crossing scheme with genotypes of parents and F_1_ males in each cross. Chromosomes derived from the ST and SR strains are shown in red and blue, respectively. **b,** Number of progeny (bar plot, mean ± S.D.; upper *X*-axis) and proportions of male progeny (circles and vertical bar indicate the proportions of each replicate and their average; lower *X*-axis) in each cross corresponding to those depicted in **a**.

The backcross using F_1_ males produced by the cross of SR females and ST males (F_1_α males, hereafter) (i.e., harboring the X chromosome from the SR strains) showed an extremely skewed female-biased sex ratio, irrespective of the female strain (crosses E and F). On the other hand, F_1_ males obtained from the cross of ST females and SR males (F_1_β males, hereafter) produced progeny with a balanced sex ratio, regardless of the female strain (crosses G and H). The difference in the extent of distortion between SR males and F_1_α males, in which the X chromosome is from the SR strain, suggested the presence of a suppressor of this X-linked driver. We hypothesized that the SR strain has an X-linked driver and a Y-linked (or autosomal recessive) “weak” suppressor, which only partially suppresses the drive, and neither is present in the ST strain. In this case, drive in the SR strain was partially suppressed, resulting in a weak female-biased sex ratio. On the other hand, drive in F_1_α males harboring the Y chromosome derived from the ST strain and heterozygous for autosomes became even more extreme, with nearly exclusive production of female offspring.

It should be mentioned that the crosses involving F_1_β males (crosses G and H) tended to produce a smaller number of progeny (<100 on average) than those of other crosses (>100 on average for all crosses, Fig. 1b; Supplementary Table S1). This observation may also be explained by the suppressor. Under this scenario, the spermatozoa in F_1_β males only contain the suppressor but not the driver, which may be deleterious for the sperm and possibly for F_1_ progeny during development due to the excess amount of the suppressor relative to the driver (or no driver).

### The X-linked driver(s) partially kills Y-chromosome-bearing sperm

To examine the effect of drive in gametes, the testis morphology and the number of sperm per cyst were compared between the SR and ST strains (Fig. 2). If the X-linked driver in the SR strain kills Y-bearing sperm, the number of sperm per cyst in SR males is expected to be smaller than that in ST males. Indeed, the average number of spermatids per cyst was significantly smaller in the ST strain than in the SR strain (124.2 ± 3.03 S.D. and 112.9 ± 9.20 S.D., *p* = 0.0104 by *t*-test; Fig. 2b). In addition, the testis of SR males was more rounded than those of ST males (Fig. 2c). However, the killing of Y-bearing sperm did not fully explain the observed sex-ratio distortion; the reduction in the number of spermatids resulted in a proportion of males of ∼45% [(112.9-124.2/2)/112.9], while the observed proportion of males was only 12–15%. Therefore, the sex-ratio distortion in the SR strain was likely caused not only by killing but also the dysfunction of Y-bearing sperm.

**Fig. 2.**
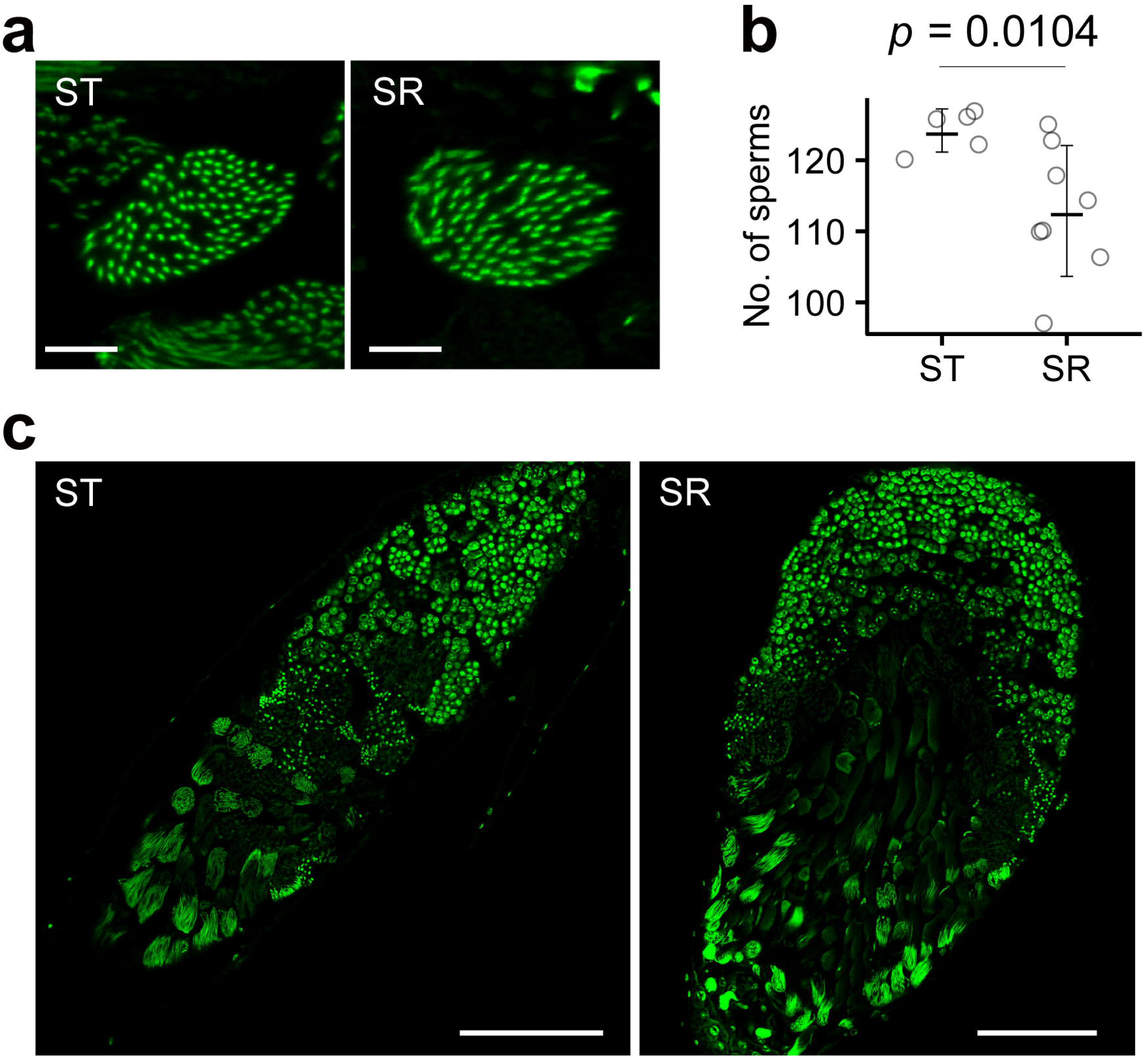
Differences in testes and cysts between the SR and ST strains. **a**, A cyst with sperm heads in the ST (left) and SR (right) strains. **b**, Number of sperm heads per cyst in the ST and SR strains; the *p*-value for comparisons between strains was calculated using the *t*-test. **c**, Whole testes of the ST (left) and SR (right) strains. Scale bars represent 5 and 100 μm in **a** and **c**, respectively.

### *Gcna* was detected as a strong X-linked candidate driver

To explore the driver on the X chromosome of the SR strain, we sequenced and assembled the male genome of the SR strain using long-read sequences generated using the Oxford Nanopore MinION and short-read sequences generated using the Illumina HiSeq X platform. We then conducted heterochromatin-enriched assembly^11^, which is particularly effective in assembling Y-derived sequences. The male genome of the ST strain was also assembled for comparison. Statistics of the final genome assemblies for the two strains are shown in Supplementary Table S2. The genome annotation obtained using the MAKER2 pipeline^12^ identified 13,119 and 12,283 genes in the genomes of the SR and ST males, respectively (Supplementary Tables S3, S4). Based on reciprocal BLASTP searches, 11,551 and 10,907 genes in the SR and ST strains were homologous with genes in *D. melanogaster*, respectively. The genes showing no homology to *D. melanogaster* sequences were assigned to 2,093 clusters using CD-HIT^13^. The SR and ST strains shared 601 clusters containing 737 and 684 genes, respectively, whereas the numbers of the clusters unique to the SR and ST strains were 796 (857 genes) and 696 (713 genes), respectively.

Because gene duplication can theoretically generate meiotic drivers^14^, duplicated genes specific to the SR genome are plausible candidates as drivers. According to the genome annotation, the duplicated genes or clusters specific to the SR strain tended to be X-linked, and the copy numbers of five of these genes or clusters were more than 10-fold higher in the SR strain than in the ST strain (Fig. 3a, Supplementary Tables S3, S4). The copy numbers of the five genes/clusters, *Pif1A* (*PFTAIRE-interacting factor 1A*), Cls. 11, *CG3726*, *Gcna* (*Germ cell nuclear acidic peptidase*), and *CG31010,* were 45, 17, 15, 14, and 13 in the genome assembly of the SR strain, respectively, but only one in the ST genome. *Pif1A* showed the largest copy number difference between the strains; however, only two were X-linked in the SR genome. Regarding Cls. 11, none of the copies were located on the X chromosome. The remaining three genes with an SR/ST copy number ratio of >10 had 4–12 X-linked copies.

**Fig. 3.**
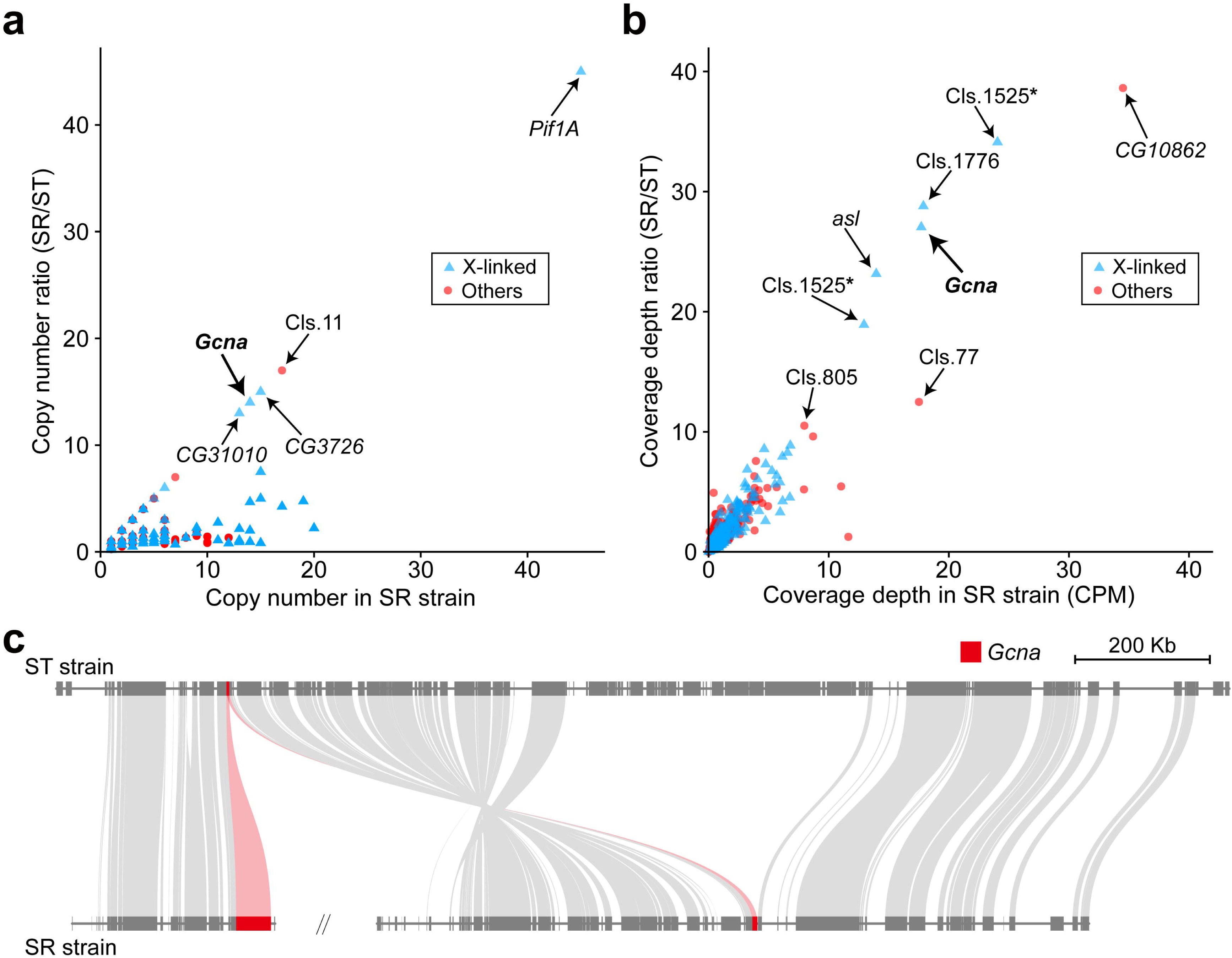
Identification of the candidate driver *Gcna* in the SR genome. **a, b,** Comparison of the copy number (**a**) and coverage depth (**b**) of each gene between the SR and ST strains, respectively. Gene/cluster names with arrows indicate the genes that showed the SR/ST ratio in terms of the copy number (**a**) or coverage depth (**b**) greater than 10. Two loci for Cls. 1525 in **b** were identified in the ST genome. Therefore, the coverage depth in the SR strain and coverage depth ratio were computed separately for each locus (indicated by asterisks). **c**, Genomic region of the *Gcna* locus and its surrounding sequence in the ST strain and its homologous region in the SR strain. Genes are indicated by thick gray blocks. To show synteny, homologous genes are connected by light gray curves. Gene synteny was visualized using JCVI toolkit^35^.

Nevertheless, the copy numbers of genes and clusters can be biased depending on the quality of the genome assembly. We therefore employed a coverage-based method as a complementary approach to examine the extent of gene duplication. Specifically, the DNA-seq short-reads from the SR and ST strains were mapped onto the ST genome, and the coverage depth was compared between the strains. The regions specifically duplicated in the SR strain are expected to show a high SR/ST coverage depth ratio. The X-linked genes or clusters with ratios of >10 were Cls. 1525, Cls. 1776, *Gcna*, and *asl* (*asterless*) (Fig. 3b, Supplementary Tables S3, S4). *Gcna* was the only gene/cluster to satisfy both criteria. In addition, *Gcna* was likely located at the edge of an inversion between the strains (Fig. 3c), consistent with previous results showing that meiotic drivers are often located within chromosomal inversions^2,3,15^. Therefore, *Gcna* was regarded as a strong candidate driver responsible for sex-ratio distortion in the SR strain.

### SprT-like domain in Gcna was massively duplicated in the SR strain and other strains with female-biased sex ratios

Gcna consists of an intrinsically disordered region (IDR) and an SprT-like metalloproteinase domain, and this domain structure is well-conserved among eukaryotes^16^. However, the SR strain showed a high coverage depth only in the 3′-half of *Gcna* encoding the SprT-like domain (Fig. 4a), raising the possibility that this domain is involved in meiotic drive. Indeed, 12 out of 14 *Gcna* loci encoded the SprT-like domain in the SR strain and many were tandemly duplicated, with 46 total copies predicted using InterProScan5^17^ (Fig. 4b). In particular, the longest, which was likely the original locus based on the genomic location (Fig. 3c), encoded as many as 21 copies of the SprT-like domain (*obs_SR_12540* in Fig. 4b). By contrast, the domain structure of Gcna in the ST strain was conserved relative to those in other eukaryotes and contained one each of the IDR and SprT-like domain (Fig. 4b). Targeted qPCR experiments further showed that the copy numbers of the genomic region encoding the SprT-like domain in the SR strain were 91.0 ± 5.3 and 154.7 ± 7.7 in males and females, respectively, when the copy number in ST females was assumed to be two (Fig. 4c), confirming the massive amplification of the region encoding the SprT-like domain on the X chromosome in the SR strain. It should be mentioned that the copy number of the region encoding the SprT-like domain on the annotated male SR genome (46 copies) was much lower than that estimated in the qPCR experiments (91 copies), implying that additional unannotated copies encoding this domain are present in the SR genome.

**Fig. 4.**
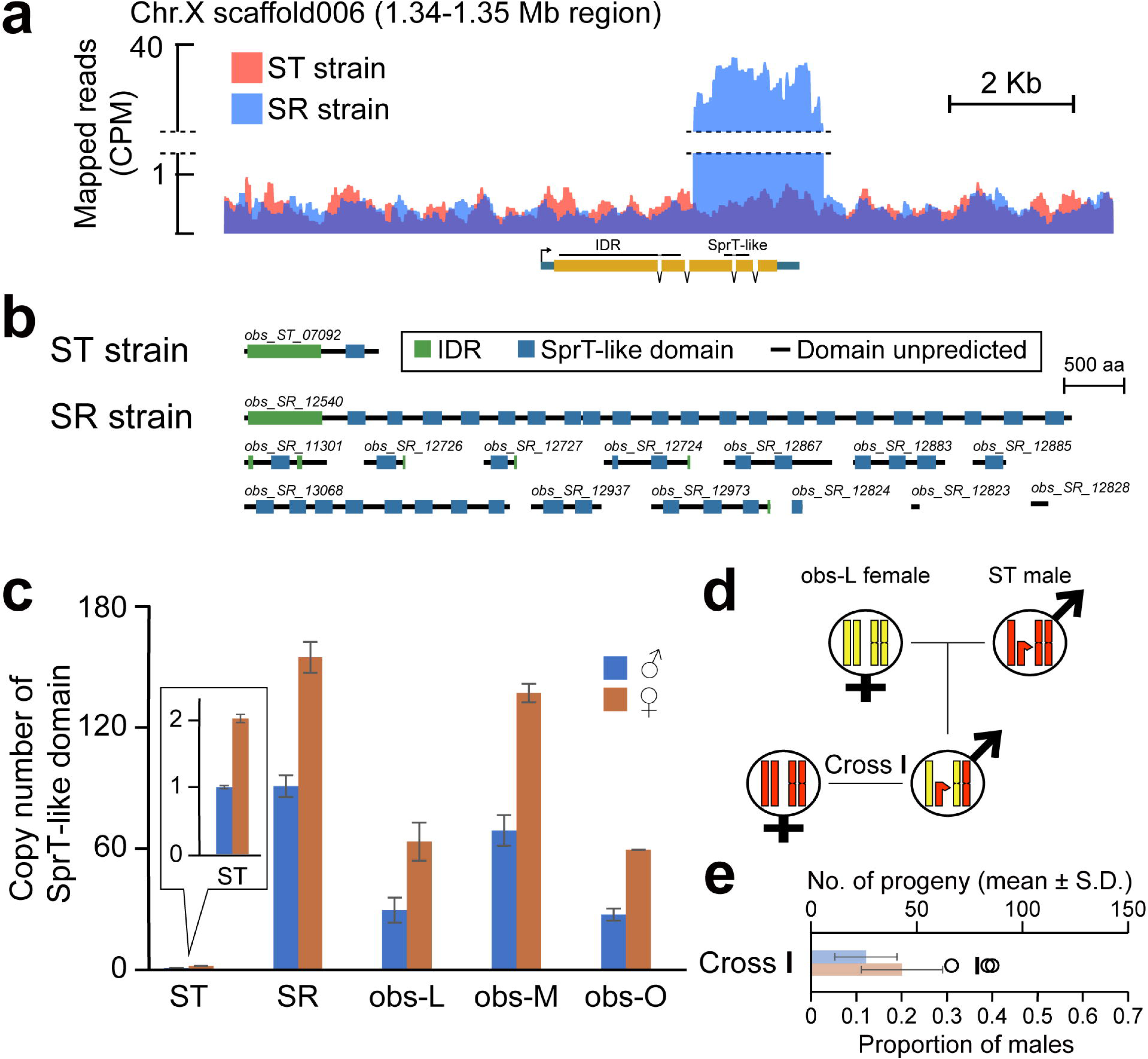
Duplication of genomic regions coding SprT-like domains in the SR and other strains but not in the ST strain. **a**, Coverage track for a part of the X chromosomal region surrounding *Gcna* in the ST genome. Gene structure of *Gcna* and its location are shown below the track (yellow: exon; light blue: UTR), with the thin black lines indicating the CDS regions coding IDR and the SprT-like domain. **b**, Protein domain structures of all *Gcna* copies predicted using InterProScan for each strain (green: IDR; blue: SprT-like domain; black line: amino acids without any known domain). IDs along the structure correspond to those in Supplementary Tables S3 and S4. **c,** Relative copy number of the SprT-like domain. The copy number in the ST males was regarded as one. **d,** Backcrossing scheme between the ST and the obs-L strains. Chromosomes derived from the ST and obs-L strains are shown in red and yellow, respectively. **e,** Number of progeny (bar plot, mean ± S.D.; upper *X*-axis) and proportions of male progeny (circles and vertical bar indicate the proportions of each replicate and their average; lower *X*-axis) in the backcross (cross I in **d**).

We further examined whether the SprT-like domain is responsible for the distortion using three other strains of *D. obscura*, obs-L, obs-M, and obs-O, with different extents of female bias^18^. Specifically, obs-L showed a balanced sex ratio, obs-O exhibited a moderate female-biased sex ratio, and obs-M showed substantial sex-ratio distortion towards female^18^. Consistent with these phenotypes, the copy number of the region encoding the SprT-like domain was highly amplified in the obs-M strain (69.1 ± 7.6 and 137.0 ± 4.6 copies for males and females, respectively), similar to results of the SR strain (Fig. 4c). The obs-O strain also showed amplification of the region encoding the SprT-like domain, but to a lesser extent, consistent with the moderate but weak distortion in this strain. Similar amplification was also observed in the obs-L strain with a balanced sex ratio, suggesting that the suppressor is strong enough to mitigate the effect of the driver in this strain. In line with this idea, the backcross experiment between males from a cross of obs-L females and ST males and ST females showed a moderate female-biased sex ratio. This observation appears reasonable because F_1_ males had the obs-L-derived X chromosome on which the genomic region encoding the SprT-like domain was moderately amplified and the ST-derived Y chromosome (and autosomes in a heterozygous condition) on which no suppressor was considered to be present (Fig. 4d,e, Supplementary Table S1).

To further examine *Gcna* as a candidate meiotic driver, RNA-seq reads from each strain were mapped to the ST genome assembly, and read counts surrounding *Gcna* were compared between strains. In females and larval and pupal males, read counts surrounding *Gcna* were higher for the ST strain than for the SR strain, although the expression level of *Gcna* was generally low. This appears reasonable because the reads were mapped to the ST genome (Fig. 5). The trend was essentially the same in adult males and testes, except in the region encoding the SprT-like domain and its surrounding regions, where the read counts were much higher for the SR strain than for ST strain (Fig. 5). These results also support our hypothesis that duplicated SprT-like domains of Gcna constitute the sex-ratio meiotic driver.

**Fig. 5.**
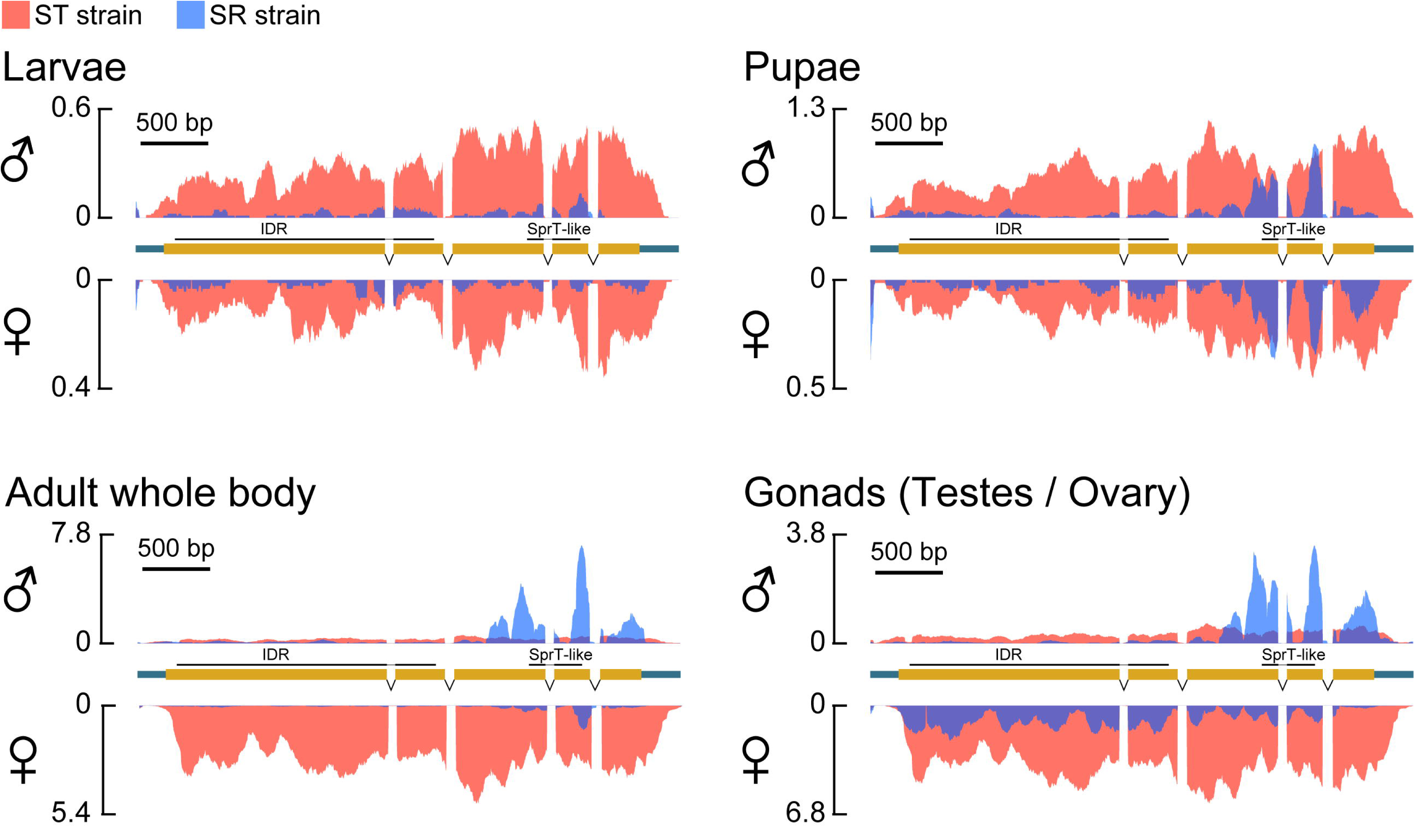
Expression levels of *Gcna* in SR and ST strains. Coverage tracks of mapped RNA-seq reads for the gene region of *Gcna* in the ST genome (above, male; below, female with flipped *Y*-axis), from the samples of larvae, pupae, adults, and gonads (testes and ovaries). Gene structure of *Gcna* is shown between the tracks of males and females.

## *miR-9c* was highly expressed in the testis of the SR strain and interacted with *Gcna*

MicroRNAs (miRNAs) provide additional layers of gene regulation at the post-transcriptional level^19^, raising the possibility that they might function as suppressors of meiotic drive. Therefore, to explore which miRNAs interact with *Gcna,* we performed small RNA-seq using adult male testis, in which meiosis occurs, of the SR and ST strains. We first searched for differentially expressed miRNAs (DEMs) among known *Drosophila* miRNAs deposited in miRBase v22^20^. We found 16 DEMs in the testis (Fig. 6a; Supplementary Table S5) and five DEMs in the male whole-body samples (Fig. 6b; Supplementary Table S6). Among these DEMs, *miR-9c* and *miR-989* showed high expression in the testis and were upregulated in the SR strain (Fig. 6a).

**Fig. 6.**
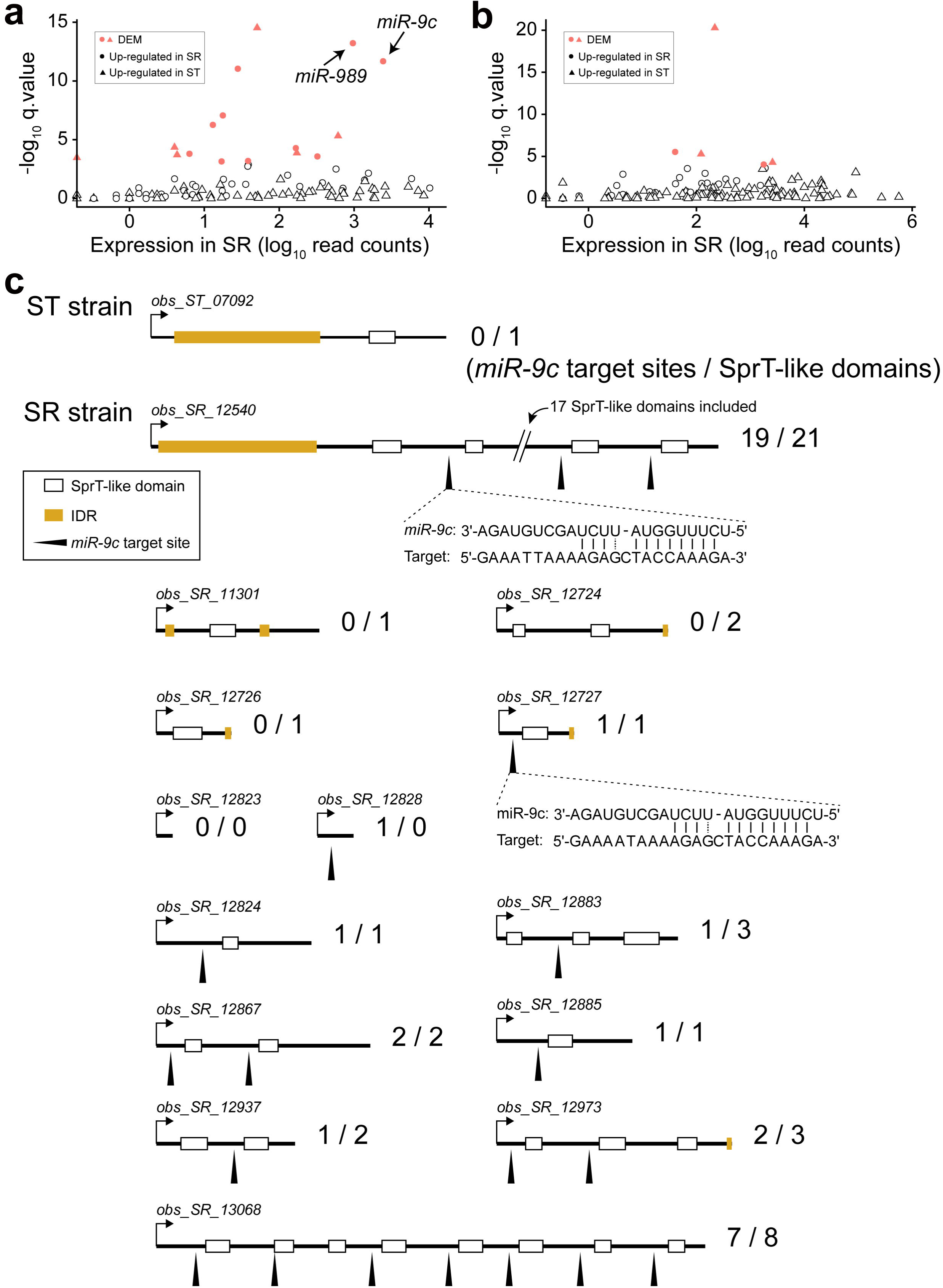
Identification of known miRNAs as candidate suppressors of *Gcna*. **a, b,** Differentially expressed known miRNAs (DEMs) in the testis (**a**) and male whole body (**b**). DEMs are indicated in red, and higher expression levels in the SR and ST strains are indicated by circles and triangles, respectively. **c,** Domain structure of Gcna and the interaction with *miR-9c* in the SR and ST strains. SprT-like domains and IDRs are indicated by white and yellow rectangles, respectively. Arrowheads indicate the predicted target sites of *miR-9c* on *Gcna*. Sequence complementarity between *miR-9c* and *Gcna* are shown below the arrowheads. The sequence complementarity was exactly the same for all loci as shown below *obs_SR_12540* except for *obs_SR_12727*. IDs along the structure correspond to those in Supplementary Tables S3 and S4.

According to miRNA target prediction, *miR-9c* was predicted to interact with 11 of 14 duplicated *Gcna* loci in the SR genome, but not to interact with the single-copy *Gcna* in the ST genome (Fig. 6c; Supplementary Table S7). Consistent with our hypothesis, the target sites of *miR-9c* on the *Gcna* copies in the SR strain tended to be located at the 5′ upstream regions of the SprT-like domains, where *miR-9c* interacted with 36 sites of, though not directly on, the 46 copies in the SprT-like domains (Fig. 6c). In addition, *miR-9c* is located on Muller B (homologous to chromosome 2L in *D. melanogaster*) in both SR and ST strains, further indicating that it functions as an autosomal suppressor of *Gcna*.

We also considered the possibility that novel miRNAs specific to *D. obscura* function as suppressors of sex-ratio distortion. Using miRDeep2^21^, we identified 456 novel miRNAs, including 51 and 54 miRNAs unique to SR and ST strains, respectively (Supplementary Table S8). Among 51 miRNAs uniquely predicted in the SR strain, 18 interacted with *Gcna*, a larger proportion than that predicted for miRNAs specific to the ST strain (11 out of 54), although the difference was not statistically significant (*p* = 0.1259 by Fisher’s exact test, Supplementary Table S9). Among the remaining novel miRNAs that were shared between the SR and ST strains, two miRNAs, *dob_miR-nov142* and *dob_miR-nov063*, showed higher testis expression in the SR strain than in the ST strain (Supplementary Table S10, Fig. S2a); in male whole-body samples, only the latter showed the same trend (Supplementary Table S11, Fig. S2b). It should be mentioned that the genomic location of *dob_miR-nov142* overlapped with *miR-9c* and the mature sequences were nearly identical, enabling both to target *Gcna*. However, the former precursor sequence was slightly shorter than the latter (Fig. S2c). Therefore, *dob_miR-nov142* may be a variant of *miR-9c* and a candidate suppressor on autosomes.

## Discussion

In this study, we investigated the genetic basis of the sex-ratio distortion in *D. obscura*, a phenomenon first reported nearly a century ago. The results can be summarized as follows: 1) The X-linked gene *Gcna* was identified as a candidate sex-ratio meiotic driver in the SR strain of *D. obscura*. 2) Meiotic drive was likely caused by massive amplification of the region encoding the SprT-like domain in *Gcna*. 3) The amplification of the SprT-like domain was a common feature of *D. obscura* strains with female-biased sex ratios. 4) *miR-9c* and several candidate miRNAs that mitigated the sex-ratio distortion were identified as suppressors. 5) The number of sperm in the SR strain was significantly lower than that in the ST strain, partly explaining the female bias. Based on these findings, we here discuss the molecular basis and evolutionary processes of the distortion, with an emphasis on the applicability of our findings to other organisms exhibiting sex-ratio meiotic drive.

The strong candidate driver, *Gcna* (*<u>g</u>erm <u>c</u>ell <u>n</u>uclear <u>a</u>cidic peptidase*), a germ cell marker in mammals^22^, is important for several processes involved in the maintenance of genome integrity, including replication, repair, transcription, and recombination in germ cells^16^. During DNA replication, DNA supercoils must be resolved by Topoisomerase II (TOP2). TOP2 interacts covalently with DNA to form transient DNA-protein crosslinks (TOP2-DPC), which are essential for genome integrity, including replication^23^. However, a fraction of the TOP2-DPC complexes are not transient and persist, resulting in the blockage of DNA replication and chromatin remodeling, which can eventually lead to DNA double-strand breaks, chromosomal segregation failure, and/or cell death^24–26^. Using the protease activity encoded by its SprT-like domain, Gcna removes these potentially deleterious persistent TOP2-DPCs^27^ (Fig. 7a).

**Fig. 7.**
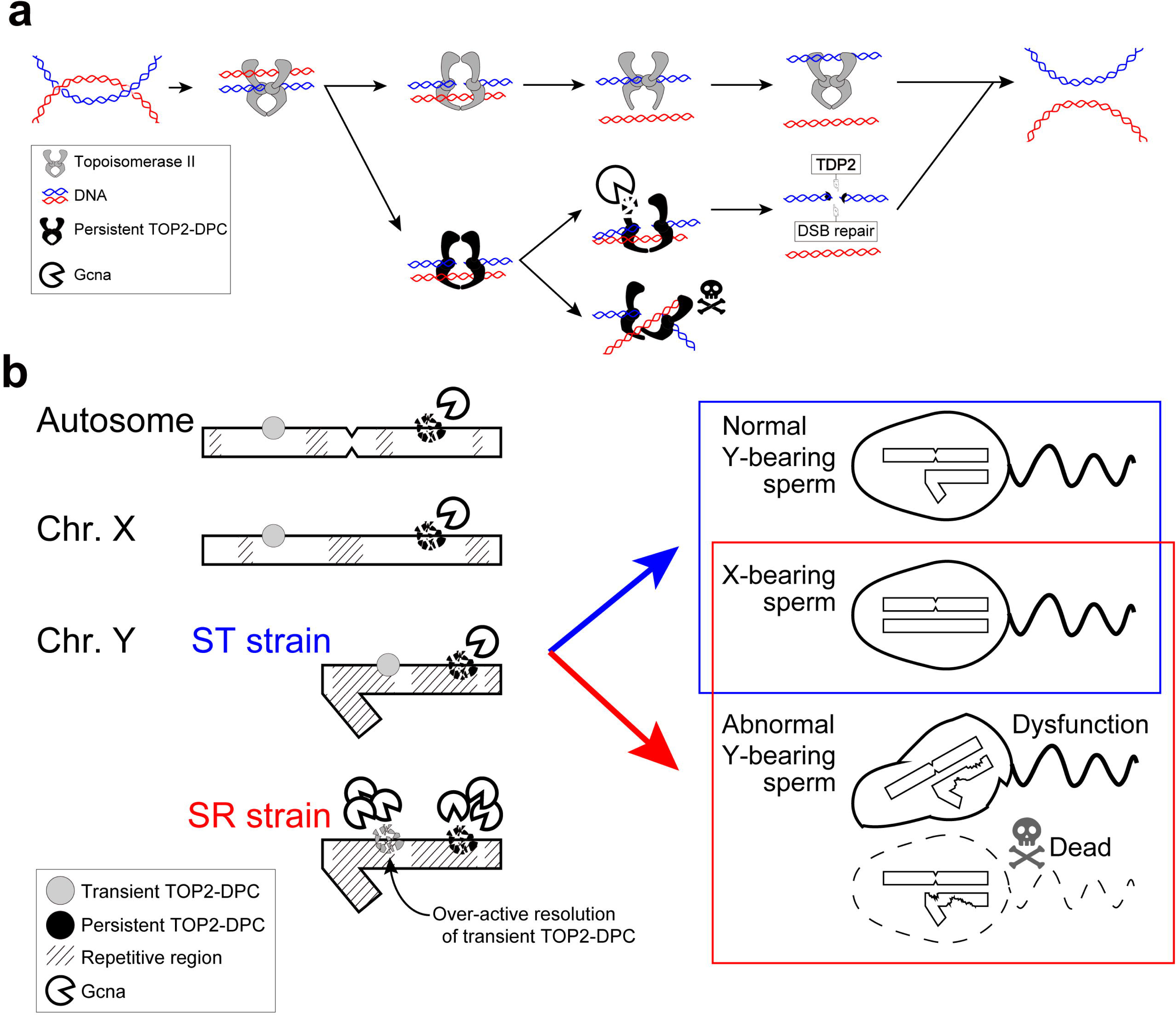
A plausible mechanism by which the driver *Gcna* causes sex-ratio distortion. **a,** Resolving DNA super coiling by transient TOP2-DPCs (above) and digestion of deleterious persistent TOP2-DPCs by *Gcna* (below). **b,** *Gcna* over-activation resulting in the digestion of both transient and persistent TOP2-DPCs on the Y chromosome. In the SR strain, the duplicated SprT-like domains likely target not only persistent but also transient TOP2-DPC on the repeat-rich Y chromosome, resulting in the killing or dysfunction of Y-bearing sperm. By contrast, the SprT-like domains target only persistent TOP2-DPC on autosomes and the X chromosome in both strains, as well as the Y chromosome in the ST strain.

We hypothesized that in the SR strain, mis-regulation or over-activation of the SprT-like domain in Gcna due to its massive amplification causes Y chromosome-specific or -biased DNA damage that kills or disrupts the Y-bearing sperm in particular during the meiotic phase of spermatogenesis, resulting in the preferred transmission of the X chromosome. As mentioned, Gcna normally digests persistent TOP2-DPCs in a specific manner. However, Gcna copies with amplified SprT-like domains in the SR strain may also digest transient TOP2-DPCs, which are essential for genome integrity. SprT-like protease activity is known to be stimulated by palindromic DNA structures frequently found in repeat-rich sequences, which are enriched in the Y chromosome^28^. Indeed, the expression level of *Gcna* in the SR strain was high in testes and adult males, particularly surrounding the region encoding the SprT-like domain. Thus, it is reasonable to postulate that the female-biased sex-ratio distortion in the SR strain is caused by targeting of Gcna with high SprT-like protease activity to TOP2-DPCs enriched on the Y chromosome (Fig. 7b).

This hypothesis is also consistent with our observation that the number of sperm cells within each cyst was significantly lower in the SR strain than the ST strain. Yet, the reduction in the number of sperm cells was less pronounced than we had expected that the number of spermatids in the SR strain would become about 40% less than that in the ST strain. The damage caused by the over-active digestion of TOP2-DPCs may depend on the frequency and locations of TOP2-DPC formation, the level of the level of SprT-like protease activity, and the expression level of *miR-9c*, which may result in varying degrees of damage, i.e., cell death, dysfunction, or no change (Fig. 7b). Tracing spermatogenesis in more detail might clarify further the mechanism of action responsible for this sex-ratio meiotic drive.

The SprT-like domain is also reported to target the X chromosome, which likely causes hybrid incompatibility^29^. The gene *maternal haploid* (*mh*) also encodes the SprT-like domain and targets particular types of satellite repeats. In *D. simulans*, the X chromosome is not targeted because it does not contain the target repeat sequences. On the other hand, target satellite repeats are present on the X chromosome in *D. melanogaster*. However, the X chromosome is not targeted in this species either, because the Spartan protease activity encoded by *mh* in *D. melanogaster* (*mh*^mel^) is very weak. When *mh*^mel^ is replaced with its homolog in *D. simulans* (*mh*^sim^), however, mh^sim^ with normal SprT-like domain activity targets not only the persistent but also the transient TOP2-DPCs surrounding satellite repeats on the X chromosome, resulting in ovarian cell death, reduced ovary sizes, and the loss of mature eggs^29^. The balance between satellite DNAs and Spartan protease activity has been proposed to be an example of antagonistic coevolution^30^. In this model, the excessive Spartan protease activity involved in TOP2-DPC degradation inactivates the Y-bearing sperm, resulting in a female-biased sex ratio. Our findings are consistent with this model.

We identified another type of possible antagonistic coevolution—the driver and suppressor. To suppress *Gcna* with hyper-protease activity possibly driven by the duplicated SprT-like domain (Fig. 7b), several miRNAs likely target and inactivate many of the duplicated domains. In particular, *miR-9c* and its possible variant, *dob_miR-nov142*, are strong candidate suppressors. These miRNAs are almost identical in their mature sequences (Supplementary Fig. S2c) and are strongly expressed in the SR testis and weakly expressed in the ST testis. In addition, these miRNAs do not target *Gcna* of the ST strain. Moreover, some of the *Gcna* loci with duplicated SprT-like domains likely escape suppression by these miRNAs due to insufficient sequence complementarity, indicating that the effect of the driver is likely stronger than that of the miRNA suppressors, explaining why the sex ratio is still biased toward females in the SR strain. The model of antagonistic coevolution between the driver and suppressor in cases of meiotic drive has already been proposed^31,32^. Indeed, in the Winters SR system of *D. simulans*^10,33^, both the driver gene *Dox* and suppressor locus *Nmy* (*Not much yang*) showed signatures of positive selection due to the biased transmission of the driver and selection on the suppressor for antagonizing the skewed sex ratio, suggesting an evolutionary “arms race” between drivers and suppressors^34^. However, the molecular bases of the interactions and their effects on meiotic drive have remained unclear. To our knowledge, this study is the first to identify a candidate driver and suppressors and to disentangle the molecular mechanism responsible for sex-chromosome drive. Strictly speaking, however, the candidate suppressors identified are located on autosomes, and it is not clear whether suppressors are also located on the Y chromosome. We intend to address this by improving the genome assemblies of SR and ST males with a special focus on the Y chromosome.

As mentioned, sex-chromosome drive resulting in a female-biased sex ratio was first documented in *D. obscura*^1^, but has remained uninvestigated for nearly 100 years. It is unfortunate that the strains used in this study are different from those used in the original study^1^. However, both the SR strain collected in Germany and the obs-M strain originating in Serbia^18^ harbored a large number of *Gcna* loci with amplification of the SprT-like domain, indicating that a shared molecular mechanism caused the female-biased sex-ratio, involving the hyper-protease activity conferred by the SprT-like domain. This study therefore resolves a long-standing question regarding sex-ratio distortion and provides guidance for addressing perennial issues in classical genetics and evolutionary biology using genomic approaches.

### Data availability

All sequence data newly generated in this study were deposited to DDBJ (accession numbers XXXXX-YYYYY). Genome assemblies obtained in this study were also deposited to DDBJ (accession numbers ZZZZZ-WWWWW). Detailed information is provided in Supplementary Table S12.

## Acknowledgments

We would like to express our sincere thanks to Mihailo Jelić and Pavle Erić at University of Belgrade for providing the Serbian *D. obscura* strains. We also thank Yasuko Ichikawa and Ikuko Ishikawa for helping with experiments. This work was supported by JSPS KAKENHI Grant Numbers 22H05073, 21H02539, and 25H01007 and Tokyo Metropolitan University.

## Author contributions

TKK and MN designed research; TKK, YC, NT, and MN conducted experiments; TKK analyzed data; and TKK, NT, and MN wrote the manuscript. All authors have confirmed the contents of the manuscript.

## Competing interests

The authors declare no competing interests.

## Methods

### Fly strains

The two *D. obscura* strains, the SR strain (strain code: 14011-0151.01; originated in Germany) and ST strain (obs-sk; Finland), were originally obtained from the National Drosophila Species Stock Center (https://www.drosophilaspecies.com) and Ehime-fly (https://kyotofly.kit.jp/cgi-bin/ehime/index.cgi), respectively. Three additional strains of *D. obscura*, obs-L, obs-M, and obs-O, used for the cross experiment and qPCR analyses, were obtained from Drs. Mihailo Jelić and Pavle Erić at University of Belgrade. All five strains have been maintained as living stocks at Tokyo Metropolitan University.

### Tetracycline treatment

Ten males and females of the SR strains of *D. obscura* were reared for two generations with standard fly medium containing tetracycline hydrochloride (0.025% w/v final concentration) at 20°C under full light. The control experiment was conducted following the same procedure with the addition of distilled water instead of tetracycline. Three replicates were prepared for each condition. Finally, the obtained adult flies were counted to calculate the sex ratio.

### Cross experiments

Reciprocal crosses were carried out using the SR and ST strains, and the F_1_ males produced by the inter-strain crosses were backcrossed with females of either the SR or ST strain. For each cross, five adult virgin females and males were placed in a glass vial with a standard fly medium. After 2 or 3 days of egg-laying, the flies were transferred into new vials, and this process was repeated seven to nine times in total with ∼21 days egg-laying. Female and male offspring obtained from the crosses were counted separately. Six replicates were prepared for each crossing condition. As controls, intra-strain crosses were also conducted for the SR and ST strains.

In addition, females of the obs-L strain exhibiting a balanced sex ratio^18^ were crossed with males of the ST strain showing a balanced sex ratio, and the resulting F_1_ males were backcrossed with females of the ST strain. Three replicates were prepared to consider variation. All crosses were conducted at 20°C under full light.

### Observation of testis and spermatids

Testes were isolated by dissecting adult flies of the SR and ST strains, fixed in cold methanol for 5 min, and then washed with ethanol. They were rehydrated through a graded ethanol series and washed in PBS with 0.1% Tween-20 (PBST). Sperm nuclei were stained with 5 μm SYTOX Green (S7020; ThermoFisher, Waltham, MA, USA) in PBST as described previously^36^. Stained testes were dehydrated in a graded ethanol series and mounted in methyl salicylate. Images were captured using a 40× Plan APO lambda lens on a NIKON AX confocal microscope (Nikon, Tokyo, Japan).

Sperm in each cyst was counted in each strain using NIS-Elements and ImageJ software^37^ (Nikon). Statistical analyses for comparing sperm counts between the SR and ST strains were performed using R version 3.5.2^38^.

### Short-read DNA, mRNA, and small RNA sequencing

Short-read DNA-seq and/or mRNA-seq were conducted for the SR and ST strains. For the SR strain, mRNA-seq data were retrieved from Nozawa et al.^39^ For DNA-seq, DNA was extracted from five females and males, separately, using Geno Plus™ Mini (Viogene, Salt Lake City, UT, USA), and libraries were constructed using the Collibri PCR-free ES DNA Library Prep Kit for Illumina (Invitrogen, Waltham, MA, USA). For mRNA-seq, total RNA was extracted from five individuals of third instar larvae, pupae, and adults for each sex with three replicates using the PureLink RNA Mini Kit (Invitrogen). Total RNA was also extracted from gonads (i.e., testis and ovary) from 10 to 20 adult flies with three replicates. Then, mRNA was isolated through poly-A selection using NEBNext Poly(A) mRNA Magnetic Isolation Module (NEB, Ipswich, MA, USA). mRNA-seq libraries were constructed using the NEBNext Ultra II Directional RNA Library Prep Kit for Illumina (NEB). Paired-end (PE) sequencing was conducted by the HiSeq X (Illumina) and NovaSeq 6000 platforms at Macrogen (Seoul, South Korea) or Novogene (Singapore) for DNA-seq and RNA-seq, respectively. The fastq reads were processed using Cutadapt v3.4^40^ and SolexaQA++ v3.1.7.1^41^ to remove adapter sequences and low-quality sequences, respectively. The thresholds for low-quality sequences were defined as a quality value less than 20 for reads generated using NovaSeq 6000 or 25 for HiSeq X and read length less than 50 nt.

Small RNA-seq was conducted for the ST and SR strains. Small RNA was extracted from adult males and testis in each strain using the NucleoSpin miRNA Kit (Macherey-Nagel, Düren, Germany) with three replicates. The small RNA-seq libraries were constructed using the NEBNext Multiplex Small RNA Library Prep Set (NEB) and sequenced on the NovaSeq 6000 platform. The obtained reads were processed in same way as those from mRNA-seq without using the length cutoff in SolexaQA++.

### Long-read genome sequencing

High-molecular-weight DNA was extracted from ∼40 adult males in each of the SR and ST strains using the Nanobind BIG DNA Kit (Circulomics, Baltimore, MD, USA), and libraries were constructed using the Nanopore Ligation Kit (ONT: Oxford Nanopore Technologies, Oxford, UK) and NEBNext Companion Module Kit (NEB). The sequencing was conducted using Oxford Nanopore MinION (ONT) with R9.4 flowcells, and the fast5 reads were subjected to base calling using Guppy v6.4.6. The obtained fastq reads were then processed using Porechop v0.2.4 and NanoLyse v1.2.1^42^ to remove adapter sequences and sequences derived from the lambda phage that was added during library construction as a control, respectively. NanoFilt v2.8.0^42^ was then used to trim low-quality sequences (<Q10).

### Genome assembly

We assembled the male genomes for the SR and ST strains as follows. First, the processed Oxford Nanopore MinION reads (ONT reads) for each strain were assembled using Flye v2.8.1 or v2.9^43^. The assemblies were processed using Racon^44^, Medaka^45^, and Pilon^46^. The scaffolds in each assembly were tentatively assigned to chromosomes as described in the “Chromosome assignment” section.

These assemblies were used as the “initial” genome assemblies for the heterochromatin-enriched assembly method (HE-assembly)^11^, which is particularly useful to obtain Y-chromosome assemblies. The processed ONT reads were mapped onto the initial assembly using minimap2^47,48^, and the unmapped reads as well as reads mapped to scaffolds that were not assigned to any chromosome were extracted. Then, the extracted reads were assembled *de novo* using Canu v2.2^49^ and Flye and processed as described above. Finally, the scaffolds assigned to the chromosomes in the initial genome assembly and the HE-scaffolds were merged using quickmerge^50^ to generate the final assemblies. The quality and the completeness of each assembly were evaluated using BUSCO v5.5.0^51^ with the diptera_odb10 dataset^52^.

### Chromosome assignment

The scaffolds in the genome assemblies were first assigned to chromosomes (i.e., Muller elements) by the following three methods: 1) the synteny of orthologs on the chromosomes of *D. melanogaster* for autosomes and the X chromosome, 2) the coverage depth ratio between female and male reads for sex chromosomes, and 3) Y chromosome Genome Scan (YGS)^53^ for the Y chromosome. For method 1), the amino-acid sequences of *D. melanogaster* (FlyBase version R6.41, https://flybase.org) were aligned to each of the *D. obscura* genome assemblies using tBLASTn v2.2.31+. If a scaffold contained more than ten genes homologous to *D. melanogaster* genes and ≥2/3 of these were located on the same chromosome arm, the so-called Muller element^54^, in the *D. melanogaster* genome, the scaffold was assigned to that Muller element. For method 2), the qualified DNA-seq short-reads from male and female samples were mapped separately onto each genome assembly using BWA-MEM v0.7.17 with the “-M” option. The coverage depths were normalized by CPM (count per million) using bamCoverage v3.5.1^55^, and the coverage ratios between males and females were compared using multiBigwigSummary v3.5.1^55^ in “BED-file” mode with the “--outRawCounts” option. The input bed-file specified the total length of each scaffold. The scaffolds with male-to-female depth ratios of >1.5 and <0.5 were assigned to the X and Y chromosomes, respectively. For method 3), YGS was performed following the original protocol^53^ using jellyfish v2.3.0^56^.

### Genome annotation and estimation of gene expression levels

All transcripts in each genome were predicted using the HISAT2-StringTie pipeline as follows. The qualified mRNA-seq reads from each stage or tissue were first mapped onto the genome assemblies using HISAT2^57^ with the “--downstream-transcriptome-assembly,” “--secondary,” and “--rna-strandness RF” options. The bam file obtained for each stage or tissue was then sorted by genome coordination using SAMtools v1.7^58^. With the sorted bam file, the transcripts for each stage or tissue were assembled using StringTie v2.2.1^59^ with the “--rf” and “-M 1.0” options, and the transcripts from all stages and tissues were finally merged using StringTie with the “--merge” option.

Genome annotation was performed using the MAKER2 pipeline^12^ with Augustus v3.4.0^60^ and SNAP^61^ as *ab initio* gene predictors. The amino-acid sequences of *D. melanogaster* (version R6.41), *D. pseudoobscura* (GenBank accession: GCA_009870125.2), and *D. obscura* (GenBank accession: GCA_018151105.1) were used as the “Protein Homology Evidence,”, and the “EST Evidence” were inputted as the transcriptome sequence (fasta format) and annotation (gff format) files generated using the HISAT2-StringTie pipeline as described above. We run three rounds of the MAKER2 annotation with “fly” as the Augustus species model. For the first round, SNAP D.melanogaster.hmm (HMM: Hidden Markov Model) was used as “snaphmm” with the parameters “est2genome” and “protein2genome” of 1. All gene models generated from the first round were merged using the “gff3_merge” script and converted to the ZFF format with the “maker2zff” script in MAKER2. These data were subjected to SNAP training with “fathom” and “forge,” and the HMM file for the second round was generated using the “hmm-assembler.pl” script. For the second round, the new HMM file generated by SNAP training was used as “snaphmm” and “est2genome” and “protein2genome” of 0. For the third round, the procedure was the same as that for the second round. The obtained annotation file was converted into the GFF3 format with gene IDs using the “maker_map_ids” and “map_gff_ids” scripts.

To estimate the expression level of each gene annotated by the above pipeline, the qualified mRNA-seq reads were mapped onto the predicted genes using STAR v2.7.6a^62^ and gene counts were converted into TPM (transcript per million) using RSEM v1.3.1^63^. The genes with TPM ≥ 1 were finally regarded as valid annotated genes. In BLASTP searches against *D. melanogaster* sequences, genes with an e-value of ≤ 1e-10 were regarded as homologs. Other genes were clustered based on the nucleotide homology using CD-HIT^13^ with default parameters, and the genes in each cluster were regarded as homologs. For functional annotation, all predicted genes were translated into amino-acid sequences and subjected to domain prediction analysis using InterProScan5^17^.

The expression of *Gcna* was compared between ST and SR strains. The expression levels were estimated using the STAR-RSEM pipeline as described above. Gene expression levels were compared based on the coverage track of mapped RNA-seq reads throughout the gene region, because the normal indices such as TPM and CPM cannot address domain-specific duplication in details. The coverage tracks were calculated by using bamCoverage v3.5.1^55^ with option “-bs 1” and “--normalizeUsing CPM”.

### Estimating the copy number ratio and coverage depth ratio for each gene

To compare the number of paralogs for each gene between the SR and ST strains, the SR/ST ratio for each gene was estimated based on the copy number and coverage depth. In the copy number-based approach, the ratio was computed by counting the paralogs that were homologous to each *D. melanogaster* gene and genes for each cluster (see the “Genome annotation and estimating gene expression level” section) in the SR and ST strains.

In the coverage depth-based approach, the DNA-seq short-reads for each sex of each strain were separately mapped onto the ST genome assembly using BWA-MEM v.0.7.17 with the “-M” option. Then, based on the CPM for each gene for each strain computed using bamCoverage v3.5.1^55^, the SR/ST ratio for the coverage depth for each gene was calculated using multiBigwigSummary v3.5.1^55^ in “BED-file” mode with the “--outRawCounts” option. For this analysis, only the genes that showed a CPM of ≥0.1 in either males or females for at least one strain were included to avoid biases caused by a small CPM. The coverage tracks were visualized using SparK^64^.

It should be mentioned that if at least one gene for each *D. melanogaster* gene or cluster was located on the scaffold assigned to the X chromosome in the SR strain, the gene or cluster was defined as X-linked in both approaches.

### Small RNA-seq analyses

Since PE reads were inapplicable for some small RNA analysis software, we first converted the PE small RNA-seq reads into single-end (SE) reads using fastq-join^65^ with default parameters. The SE reads were then mapped onto the known tRNA and rRNA sequences of *D. melanogaster* downloaded from RNAcentral^66^ using bowtie2 v2.4.4^67^ with the “--very-sensitive” mode, and the unmapped reads (i.e., neither tRNA nor rRNA sequences) were extracted using SAMtools. The extracted reads were then annotated using miRDeep2 software^21^. For known miRNAs, the small RNA-seq reads from testis and adult male whole-body samples for each strain were mapped and quantified using the quantifier.pl script included in miRDeep2. The output containing the read counts of each miRNA was used for detecting differentially expressed miRNAs (DEMs) using the TCC package^68^ implemented in R software v4.4.5^69^. For novel miRNA predictions, the Perl script mapper.pl, included in miRDeep2, was used for mapping. Then, the miRDeep2.pl script was run with the mapping file (arf format), small RNA reads, and known miRNAs (precursor and mature sequences) of *Drosophila* with the “-t D.pseudoobscura” option. All known miRNA sequences in *Drosophila* species were downloaded from miRBase v22^20^. The novel miRNAs predicted in each strain were then named based on sequence identity and chromosomal locations. A novel miRNA with an identical sequence and homologous genomic locations in the SR and ST genomes was regarded as an orthologous miRNA and the same name was given, whereas all other novel miRNAs were considered to be strain-specific and were given unique names.

The DEMs of known miRNAs with higher expression in the SR strain and all novel miRNAs were then subjected to target prediction. Prediction was performed using miRanda software^70^ with the “-sc 150.0” and “-en 20” options, using the mature miRNA sequences and transcript sequences of *D. obscura* generated according to the gene annotations. Considering the possibility that not only UTRs but also CDS can be targets of miRNAs in animals^71^, the entire transcript for each gene was used for target prediction.

### qPCR for detecting duplications of the SprT-like domain

The five strains of *D. obscura* (i.e., SR, ST, obs-L, obs-M, and obs-O) were used for quantitative PCR (qPCR). DNA was extracted as mentioned above. qPCR primers were designed at the conserved site among the copies of the SprT-like domain in the SR and ST genomes. *RpS3* (*Ribosomal protein S3*) was used as the reference for a single-copy autosomal gene. qPCR was performed using PowerUp SYBR Green Master Mix (Thermo Fisher) and LightCycler 96 (Roche, Basel, Switzerland) under standard conditions. The relative copy number of the SprT-like domains was estimated using the Pfaffl method^72^, and the absolute copy number of the SprT-like domains for females and males in each strain was calculated by defining the copy number in the SR females as two. The primer sequences were as follows: SprT-F 5′-TGCTGACCACAGCTGACAGAC-3′; SprT-R 5′-GGCACAGCTCGTGGATGAG-3′; RpS3-F 5′-CTCCGACGGTATCTTCAAGG-3′; and RpS3-R 5′-ACACCGGAGTAACCATCCTC-3′.

## Notes

### Competing Interest Statement

The authors have declared no competing interest.

