## Supplementary Figure S1 and S2 for "Disentangling sex-ratio meiotic drive in *Drosophila* a century after its original discovery"

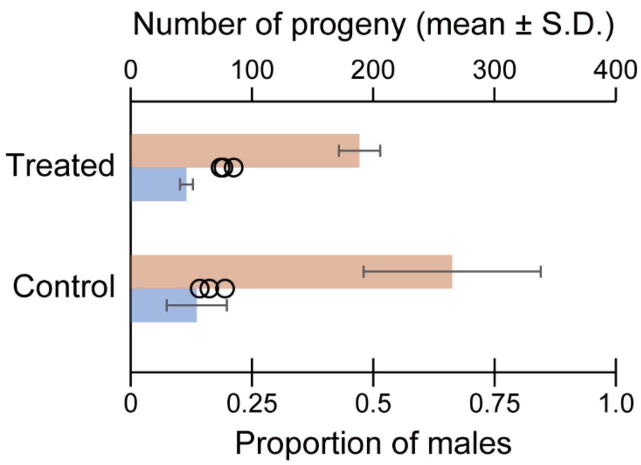


**Supplementary Fig. S1. Number of progeny (bar plot, mean ± S.D.; upper *X*-axis) and proportions of male progeny (circles and vertical bar indicate the proportions of each replicate and their average; bottom *X*-axis) in the $F_2$ generation of tetracycline-treated and control flies.**

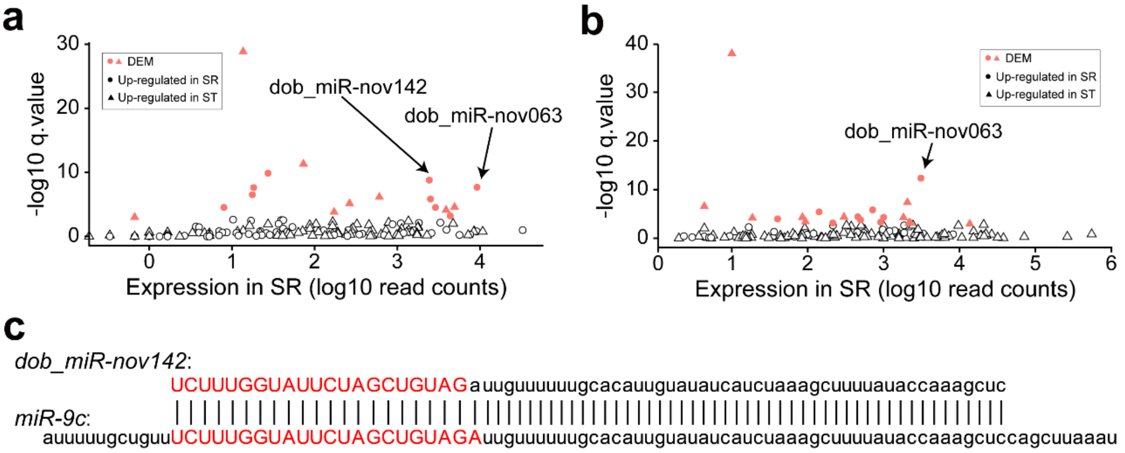


**Supplementary Fig. S2. Identification of novel miRNAs as candidate suppressors of *Gcna*. a, b,** Differentially expressed novel miRNAs (DEMs) in the testis (**a**) and male whole body (**b**). DEMs are indicated in red, and higher expression levels in the SR and ST strains are indicated by circles and triangles, respectively. **c**, Sequence alignment of the precursor regions of *miR-9c* and *dob_miR-nov142*. Nucleotides with uppercase letters in red indicate the mature sequence of these miRNAs.
